# A bidirectional interaction between the SREBP pathway and the LINC complex component nesprin-4 controls lipid metabolism

**DOI:** 10.64898/2026.04.18.719359

**Authors:** Badria Fouad Al-Sammak, Hafiz Majid Mahmood, Maria Teresa Bengoechea-Alonso, Henning F. Horn, Johan Ericsson

## Abstract

This report identifies a bidirectional signaling axis connecting lipid metabolism to nuclear mechanotransduction, with the potential to control fatty acid/triglyceride metabolism. The sterol regulatory element-binding (SREBP) family of transcription factors control fatty acid, triglyceride and cholesterol synthesis and metabolism. The family consists of three members: SREBP1a, SREBP1c, and SREBP2, that are regulated by intracellular cholesterol levels and insulin signaling. The SREBP2-dependent control of the LDL receptor gene is a well-established target for cholesterol-lowering therapeutics and the activity of SREBP1c is an attractive target in metabolic disease. In the current report, we identify SYNE4 (nesprin-4), a component of the Linker of Nucleoskeleton and Cytoskeleton (LINC) complex, as a direct target of the SREBP family of transcription factors, and show that nesprin-4 in turn supports SREBP1c function. We identify functional SREBP binding sites in the human SYNE4 promoter and demonstrate that these are required for the sterol– and SREBP-dependent regulation of the promoter. Furthermore, we show that the endogenous SYNE4 gene is also regulated by SREBP1/2 and intracellular sterol levels. Interestingly, SREBP2 is responsible for the sterol regulation of the SYNE4 gene in HepG2 cells, while SREBP1 is the major regulator in MCF7 cells, demonstrating that different cell types use different SREBP paralogs to regulate the same promoter/gene. Importantly, we find that nesprin-4 is a positive regulator of SREBP1c expression and function in HepG2 cells and during the differentiation of human adipose-derived stem cells. In summary, the current report identifies a novel regulatory interaction between lipid metabolism and the LINC complex. Importantly, we demonstrate that this signaling axis is bidirectional, forming a closed loop that has the potential to control SREBP1c activity and thereby fatty acid and triglyceride synthesis/metabolism. Based on our data, we propose that the nesprin-4-dependent regulation of SREBP1c could represent a novel therapeutic target in metabolic disease.

## Introduction

The nuclear envelope (NE) is an important barrier that protects the genome from biochemical and mechanical stress. It consists of a double lipid bilayer membrane and an underlying nuclear lamina. The outer nuclear membrane (ONM) forms a continuous membrane system with the endoplasmic reticulum. The inner nuclear membrane (INM) sits between the ONM and the nuclear lamina and is host to a number of membrane-associated proteins that confer unique functionality to the INM (recently reviewed in [1]). Underneath the INM sits the nuclear lamina, which is a meshwork of intermediate filament proteins that confer mechanical resilience to the NE (recently reviewed in [2]).

The NE integrates a host of cytosolic and nucleoplasmic signals and events. Key to this integration are transmembrane proteins that reside within the NE, including LINC (<u>Li</u>nker of <u>n</u>ucleoskeleton and <u>c</u>ytoskeleton) complex proteins. LINC complexes consist of ONM KASH domain proteins that are tethered to INM SUN domain proteins, creating a physical connection from cytoskeleton to nucleoskeleton [3–5]. LINC complexes play a central role in nuclear positioning and in mechanotransduction and have been implicated in several developmental and disease processes [6].

We recently showed that nesprin-4, a KASH domain protein primarily expressed in secretory epithelia [7], plays a role in breast cancer migration and invasion [8]. We observed that nesprin-4 expression changes significantly during tumorigenesis, with increased expression in cancer compared to normal breast tissue. We also found that during tumor evolution towards more aggressive subtypes, nesprin-4 was downregulated, with many of the most aggressive triple-negative breast cancers expressing little or no nesprin-4 [8].

Little is known about how the expression of LINC complex proteins is regulated. A few transcription factors have been identified for some LINC components [9, 10], but only one has recently been shown to regulate nesprin-4 expression in colorectal cancer. The YEATS domain-containing protein 4 (GAS41) binds to the SYNE4 promoter in a H3K27 acetylation-dependent manner [11].

Disturbances in lipid metabolism are at the core of several global health challenges, including type 2 diabetes, obesity and cardiovascular disease [12, 13]. Sterol regulatory element-binding proteins (SREBP1a, SREBP1c, and SREBP2) are transcription factors that control the expression of genes involved in cholesterol, fatty acid and triglyceride synthesis and metabolism [14–16]. SREBP1c and SREBP2 are expressed in all mammalian cells and control genes involved in fatty acid/triglyceride and cholesterol synthesis, respectively. SREBP1a is the strongest transcription factor of the family and activates most SREBP target genes. Thus, SREBP1a is expressed in rapidly proliferating cells, including cancer cells, to ensure a sufficient supply of lipids to support cell growth. All SREBPs are synthesized as large precursor proteins that are inserted into the endoplasmic reticulum (ER) membrane and need to be proteolytically cleaved to generate the active transcription factors [17, 18]. The activation of SREBP1/2 is dependent on their transport from the ER to the Golgi, a process that is regulated by intracellular cholesterol levels and insulin signaling. In the ER, the SREBP precursor proteins interact with a sterol-sensing chaperone protein known as SREBP cleavage activating protein (SCAP) [16, 19, 20]. When mammalian cells are deprived of cholesterol, SCAP escorts SREBP1/2 in COPII vesicles from the ER to the Golgi. In the Golgi, two proteases (S1P and S2P) sequentially cleave SREBP1/2, releasing the N-terminal transcriptional domains, which translocate to the nucleus and activate genes involved in fatty acid and cholesterol synthesis and uptake [18]. When the amount of cholesterol in the ER membrane exceeds a certain threshold, cholesterol binds to SCAP, and this induces the SCAP/SREBP complex to bind to ER retention proteins, known as insulin-induced gene (Insig) 1/2. The SREBP/SCAP/Insig complex is retained in the ER, and the expression of SREBP target genes drops. From a clinical perspective, the most important SREBP target gene is the LDL receptor gene [21]. This is illustrated by the fact that the SREBP-dependent induction of LDL receptor mRNA and protein in the liver is responsible for the LDL-cholesterol lowering activity of the statin family of drugs. Both the expression and activation of SREBP1c are induced in response to insulin signaling [22–26], and have been shown to regulate insulin-dependent adipocyte differentiation in vitro. Importantly, the insulin-dependent activation of SREBP1c in the liver is preserved in T2D patients, thereby contributing to the development of non-alcoholic fatty liver disease, hypertriglyceridemia and the worsening of insulin resistance [23]. Thus, mechanisms that specifically regulate the expression/function of SREBP1c are possible targets in metabolic disease.

Here we identify a bidirectional signaling axis between SREBP1/2 and the LINC complex component nesprin-4. Initially, we show that the expression of SYNE4, the gene encoding nesprin-4, is controlled by members of the SREBP family of transcription factors through functional SREBP-binding sites in its promoter. Subsequently, we demonstrate that this functional interaction impacts on SREBP1c expression and function, at least in cell types characterized by high lipogenic potential. Taken together, our results suggest that the SREBP1/2-nesprin-4-SREBP1c axis could be a target in metabolic disease.

## Results

### The SYNE4 promoter contains SREBP1/2 binding sites

We have previously demonstrated that nesprin-4 (gene symbol SYNE4) affects breast cancer migration and intravasation [8]. Using loss– and gain-of-function assays, we found that the ectopic expression of nesprin-4 in invasive MDA-MB-231 cells enhanced their migration/invasion but inhibited their intravasation. In addition, we demonstrated that the shRNA-mediated inactivation of SYNE4 in MCF7 cells inhibited their invasiveness. Taken together, these observations suggest that the expression of nesprin-4 could affect cancer metastasis. Notably, high SYNE4 expression may promote migration/invasion, while low expression could promote intravasation, EMT and the metastatic process. To identify the factors controlling SYNE4 expression, we cloned the SYNE4 promoter (defined as the non-coding sequence separating the SYNE4 and ALKBH6 gene on chromosome 19) from human MCF7 cells. To identify the factors controlling the expression of the SYNE4 gene, we cloned its promoter from human MCF7 cells. As illustrated in Fig. 1A, the core promoter corresponds to 535 base pairs (–402 – +133) with the transcriptional start site (TSS) located at +1. The full sequence of the cloned promoter is provided in Fig. S1. A visual inspection of the promoter identified four E-box elements (CAnnTG), known to function as a binding site for basic helix-loop-helix transcription factors. Interestingly, we also identified three potential sterol regulatory elements (SREs), known to function as binding sites for the sterol regulatory element-binding (SREBP) family of transcription factors. Two of these binding sites (SRE1 and SRE2) are located upstream and the third (SRE3) downstream of the TSS. Interestingly, these sequence motifs colocalize with SREBP1 binding sites identified in genome-wide ChIP-sequencing efforts by the ENCODE project (–310 – +229 in HepG2 cells), using multiple cell lines, including HepG2 and MCF7 (https://www.encodeproject.org, Fig. 1A). These data suggest that there is a potential association between SREBP1/2 and SYNE4. To test if SREBP1 was able to interact with the SYNE4 promoter, we used recombinant nuclear SREBP1a and the cloned promoter in electromobility shift assays (EMSAs). As illustrated in Figure 1B, the addition of SREBP1a to the EMSA reaction generated a distinct DNA-protein complex. To determine if endogenous SREBP1 could interact with the SYNE4 promoter, we substituted recombinant SREBP1a with nuclear extracts from HeLa cells grown in lipoprotein-deficient media to induce the activation of SREBP1/2. Addition of the nuclear extract to the SYNE4 promoter probe generated a discrete protein-DNA complex (Figure 1C). Importantly, this complex was shifted upwards when an antibody recognizing nuclear SREBP1 was added to the EMSA reaction, suggesting that the original protein-DNA complex contains nuclear SREBP1.

**Figure 1.**
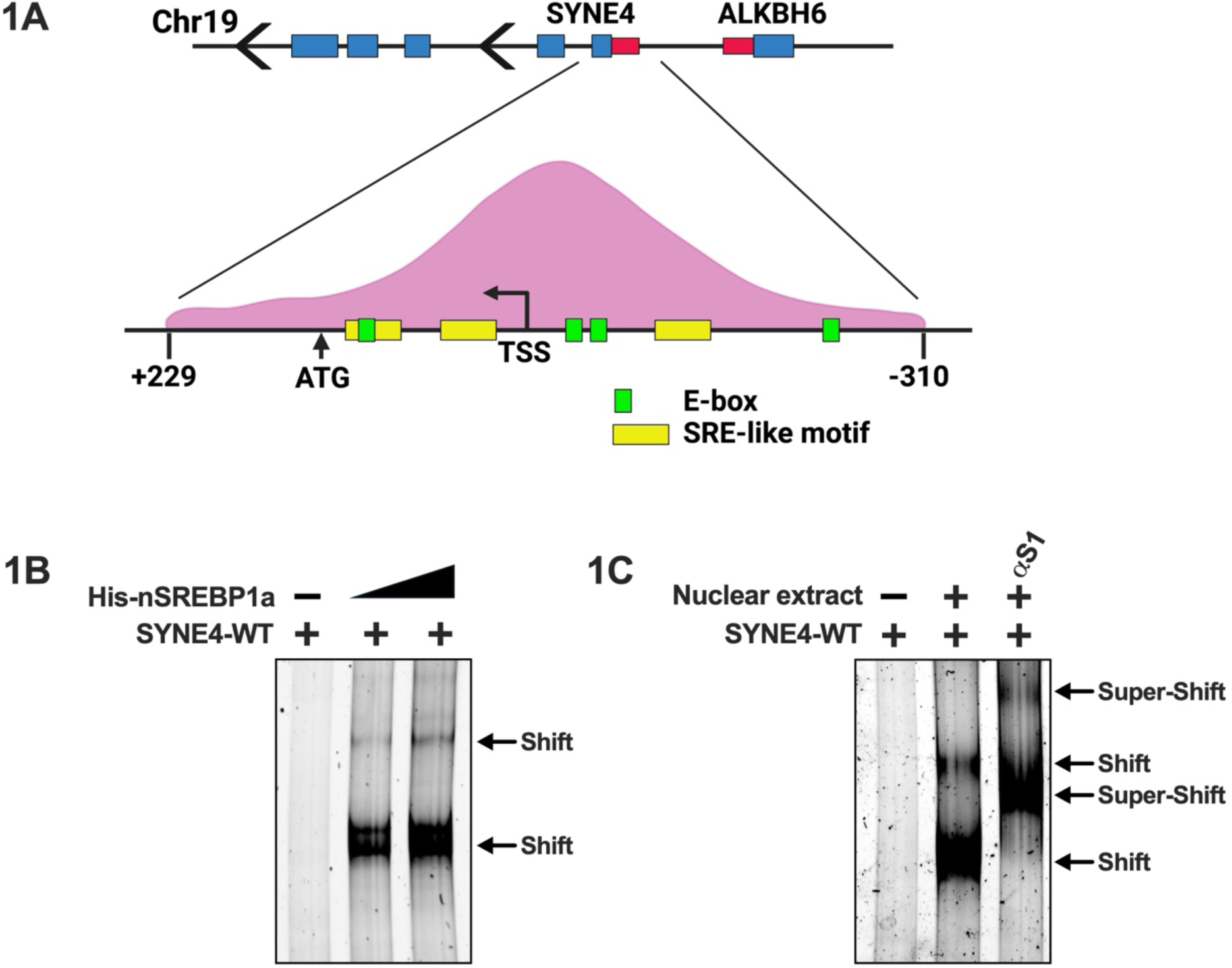
The human SYNE4 promoter contains potential SREBP binding sites. (**A**) Illustration of the human SYNE4 promoter, including the location of E-boxes, potential SREBP binding (SRE) motifs and transcriptional start site. The SREBP1 ChIP-seq peak reported by the ENCODE project is superimposed over the proximal promoter. (**B**) Increasing amounts of 6xHis-nSREBP1a were mixed with the unlabeled SYNE4 promoter probe. The reaction mixtures were resolved on native PAGE gels, and the DNA-protein complexes were visualized with SYBR Safe. The shifted DNA-protein complexes are indicated by arrows. (**C**) Nuclear extracts were prepared from HeLa cells grown in lipoprotein-deficient media and incubated with unlabeled SYNE4 promoter probe. Where indicated, SREBP1-specific antibodies were added to the reaction mixture prior to loading the samples on the gel. The shifted and antibody-specific supershifted DNA-protein complexes are indicated by arrows.

### The SYNE4 promoter is regulated by sterols and nuclear SREBP1/2

To start exploring if the SYNE4 promoter was responsive to SREBP1/2, the promoter fragment was cloned into pGL4basic. In the resulting construct, the expression of the luciferase reporter gene is under the control of the SYNE4 proximal promoter. As seen in Figure 2A, the activity of the SYNE4 promoter was relatively low in HepG2 cells grown in regular media. However, the activity was greatly induced following ectopic expression of nuclear SREBP1a (Figure 2A). Similar results were seen in cells expressing nuclear SREBP1c and SREBP2, although the induction in promoter activity was not as high as in SREBP1a expressing cells (Fig. 2A).

**Figure 2.**
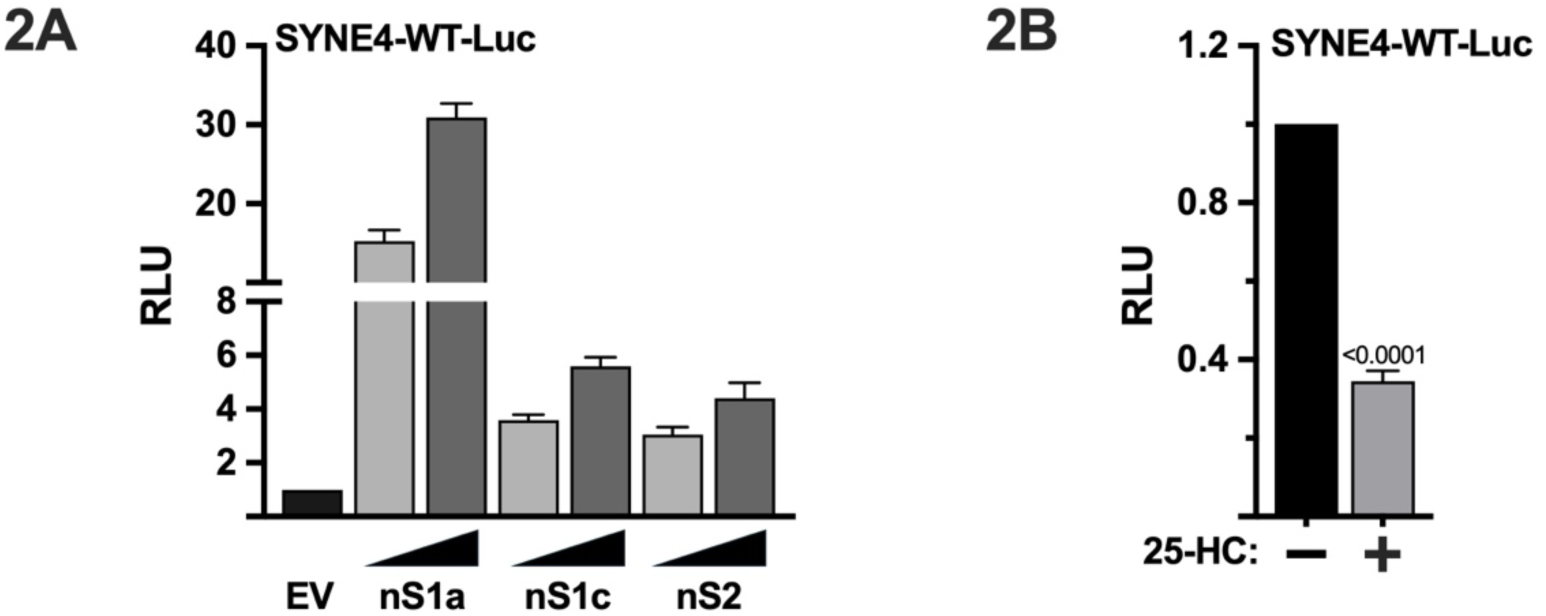
The human SYNE4 promoter is regulated by sterols and SREBP1/2. (**A**) The wild-type SYNE4 promoter-reporter construct (SYNE4-WT-Luc) was transfected into HepG2 cells together with empty vector (EV) or increasing amounts of nuclear SREBP1a (nS1a), SREBP1c (nS1c), or SREBP2 (nS2). Forty-eight hours after transfection, the cells were lysed, and luciferase activity was measured. (**B**) HepG2 cells were transfected with the wild-type SYNE4 promoter-reporter construct. Forty-eight hours following transfection, the media was replaced with lipoprotein-deficient media. Where indicated, media was supplemented with 25-hydroxycholesterol (25-HC) to block SREBP1/2 activation. Seventy-two hours after transfection, the cells were lysed, and luciferase activity was measured. The cells in A and B were also transfected with the β-galactosidase gene as an internal control for transfection efficiency. Luciferase values (relative light units, RLU) were calculated by dividing the luciferase activity by the β-galactosidase activity.

The activation of SREBP1/2 is blocked in cells grown in the presence of an excess of cholesterol and/or cholesterol metabolites, including 25-hydroxycholesterol. To test if the SYNE4 promoter was sensitive to changes in intracellular sterol levels, HepG2 cells transfected with the SYNE4 promoter-reporter construct were placed in lipoprotein-deficient media in the absence or presence of 25-hydroxycholesterol. As seen in Figure 2B, the addition of 25-hydroxycholesterol to the sterol-depleted media significantly reduced the activity of the promoter, suggesting that the SYNE4 promoter is regulated by endogenous SREBP1/2 in a sterol-dependent manner. Thus, our results show that SREBP1/2 binds to the SYNE4 promoter, and that the activity of the promoter is regulated by intracellular sterol levels and ectopically expressed nuclear SREBP1/2. To further explore the role of endogenous SREBP1/2 in the regulation of the SYNE4 promoter, we transfected HepG2 cells with the SYNE4 promoter-reporter construct in the presence of non-targeted shRNA or shRNA targeting SREBP1 or SREBP2. To induce the activation of SREBP1/2, the media was changed to lipoprotein-deficient media for the last 24 hours of the experiment. Inactivation of SREBP1 significantly reduced the activity of the SYNE4 promoter (Figure 3A), and similar results were seen following the inactivation of SREBP2. The sterol-dependent suppression of the SYNE4 promoter was retained in SREBP1-deficient cells. However, the sterol-dependent regulation of the SYNE4 promoter did not reach statistical significance in SREBP2-deficient cells, suggesting that SREBP2 is critical for the sterol-dependent regulation of the promoter in HepG2 cells.

**Figure 3.**
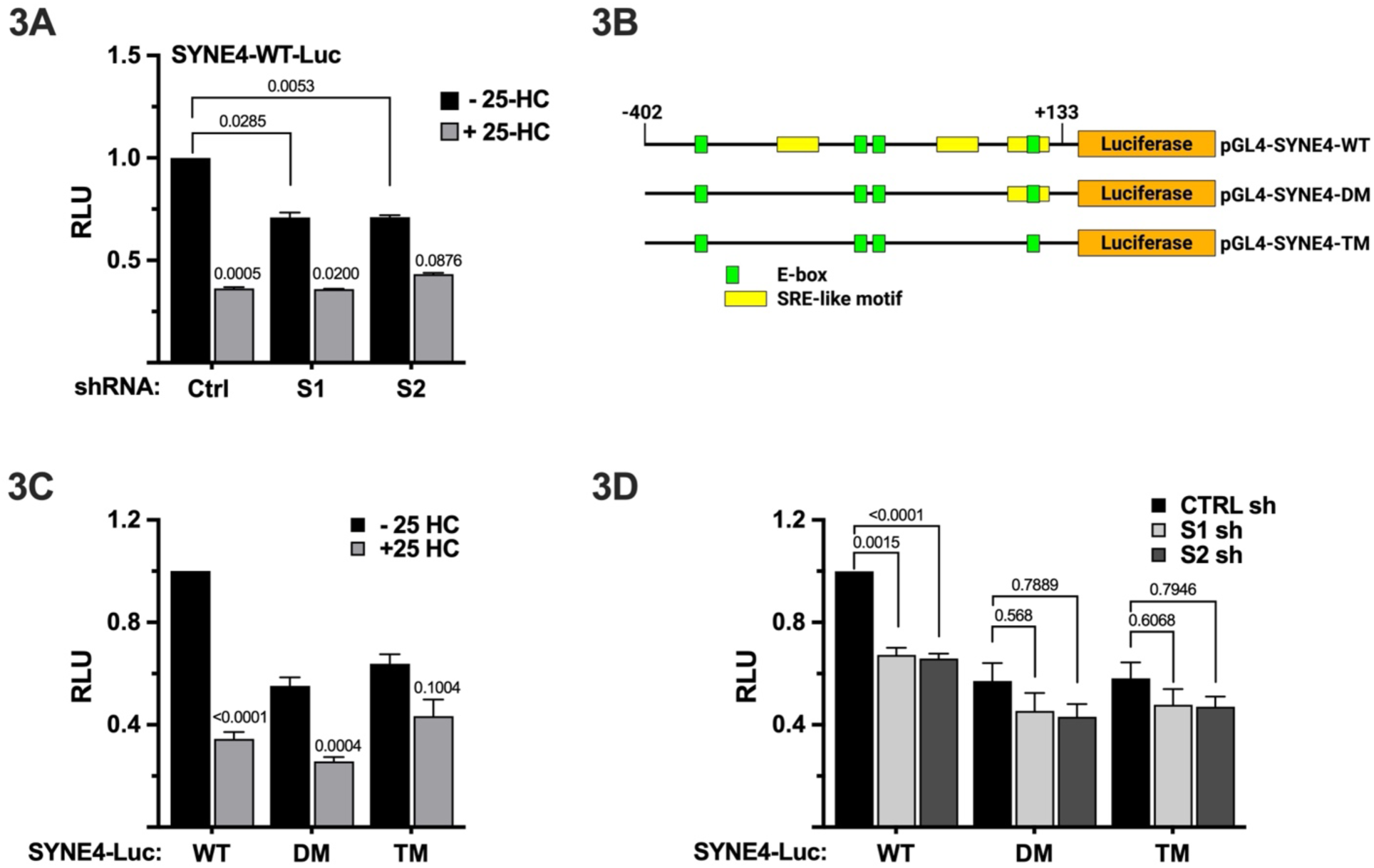
The SREBP binding sites control the sterol-sensitivity of the SYNE4 promoter. (A) HepG2 cells were transfected with the wild-type SYNE4 promoter-reporter gene (SYNE4-WT-Luc) together with non-targeted (Ctrl), SREBP1 (S1), or SREBP2 (S2) shRNA. Forty-eight hours following transfection, the media was replaced with lipoprotein-deficient media. Where indicated, media was supplemented with 25-hydroxycholesterol (25-HC) to block SREBP1/2 activation. Seventy-two hours after transfection, the cells were lysed, and luciferase activity was measured. (**B**) The wild-type and SRE-deleted versions of the SYNE4 promoter reporter gene are illustrated. DM, deletion of SRE1 and SRE2: TM, deletion of SRE1-3. (**C**) HepG2 cells were transfected with SYNE4 promoter reporter constructs, either wild-type (WT) or containing deletions of SRE1 and SRE2 (DM) or all three SREs (TM). Forty-eight hours following transfection, the media was replaced with lipoprotein-deficient media. Where indicated, media was supplemented with 25-hydroxycholesterol (25-HC) to block SREBP1/2 activation. Seventy-two hours after transfection, the cells were lysed, and luciferase activity was measured. (**D**) HepG2 cells were transfected with SYNE4 promoter reporter constructs, either wild-type (WT), DM or TM, together with non-targeted (Ctrl), SREBP1 (S1), or SREBP2 (S2) shRNA. Forty-eight hours after transfection, the media was changed lipoprotein-deficient media to induce the activation of SREBP1/2. The cells in A-C were also transfected with the β-galactosidase gene as an internal control for transfection efficiency. Luciferase values (relative light units, RLU) were calculated by dividing the luciferase activity by the β-galactosidase activity.

To determine if the sterol regulatory elements in the SYNE4 promoter were responsible for its sterol-dependent regulation, we generated promoter-reporter constructs containing deletions of the potential SRE sequences and used these constructs in promoter-reporter assays. Forty-eight hours following transfection, HepG2 cells were placed in lipoprotein-deficient media in the absence or presence of 25-hydroxycholesterol. Deleting the potential SRE motifs individually did not affect the expression or sterol sensitivity of the SYNE4 promoter. However, deleting SRE1 and SRE2 in combination (double mutant, DM) reduced both the expression and sterol-responsiveness of the promoter (Fig. 3B). Importantly, deletion of all three SRE motifs (triple mutant, TM) reduced both the activity and sterol sensitivity of the SYNE4 promoter. To determine if the SRE motifs were also responsible for the SREBP1/2-dependent regulation of the SYNE4 promoter, the wild-type, DM and TM promoter-reporter constructs used in Fig. 3B were transfected into HepG2 cells together with non-targeted or SREBP1/2 shRNA. As seen in Fig. 3C, inactivation of both SREBP1 and SREBP2 reduced the activity of the wild-type promoter. In contrast, the DM and TM mutants were insensitive to the loss of both SREBP1 and SREBP2. Taken together, these results suggest that the sterol/SREBP-dependent regulation of the SYNE4 promoter is dependent on its potential SRE motifs. Although the activity of the SYNE4 promoter is reduced in response to both SREBP1 and SREBP2 loss, it appears that SREBP2 plays a dominant role in its the sterol-dependent regulation of SYNE4, at least in HepG2 cells.

### The expression of the SYNE4 gene is regulated by sterols and SREBP1/2

The previous experiments demonstrated that a SYNE4 promoter construct is sensitive to nuclear SREBP1/2 and 25-hydroxycholesterol, an inhibitor of SREBP1/2 activation. To test if the expression of the endogenous SYNE4 gene is also sensitive to changes in SREBP1/2 activity, HepG2 cells were transduced with lentiviruses expressing non-targeted, SREBP1 or SREBP2 shRNA. The cells were grown in regular media for 72 hours before lysis and RNA extraction. As seen in Fig. 4A, the expression of SYNE4 mRNA was significantly reduced in SREBP1 and SREBP2 deficient cells, supporting the results of our promoter-reporter assays. To determine if the SYNE4 gene was sterol-sensitive and if so, whether this was dependent on SREBP1 and/or SREBP2, HepG2 cells were transduced as described above, and the cells were placed in lipoprotein-deficient media in the absence or presence of 25-hydroxycholesterol 24 hours before lysis and RNA extraction. The expression of SYNE4 was reduced in response to 25-hydroxycholesterol in cells expressing non-targeted shRNA (Fig. 4B). Inactivation of SREBP1 did not affect the sterol sensitivity of the SYNE4 gene. However, 25-hydroxycholesterol failed to reduce the expression of SYNE4 in SREBP2-deficient HepG2 cells (Fig. 4B), suggesting that SREBP2 is responsible for the sterol-dependent regulation of SYNE4 in these cells, supporting the data reported in Fig. 3A. This hypothesis was further supported when we analyzed the SREBP2 sensitivity of the wild-type and SRE-deleted (DM and TM) versions of the SYNE4 promoter in HepG2 cells. As seen earlier, the wild-type promoter was sensitive to increasing levels of nuclear SREBP2 (Fig. 4C). However, the SREBP2 responsiveness of both the DM and TM promoters was significantly reduced, supporting the notion that SREBP2 is an important regulator of the SYNE4 promoter in HepG2 cells.

**Figure 4.**
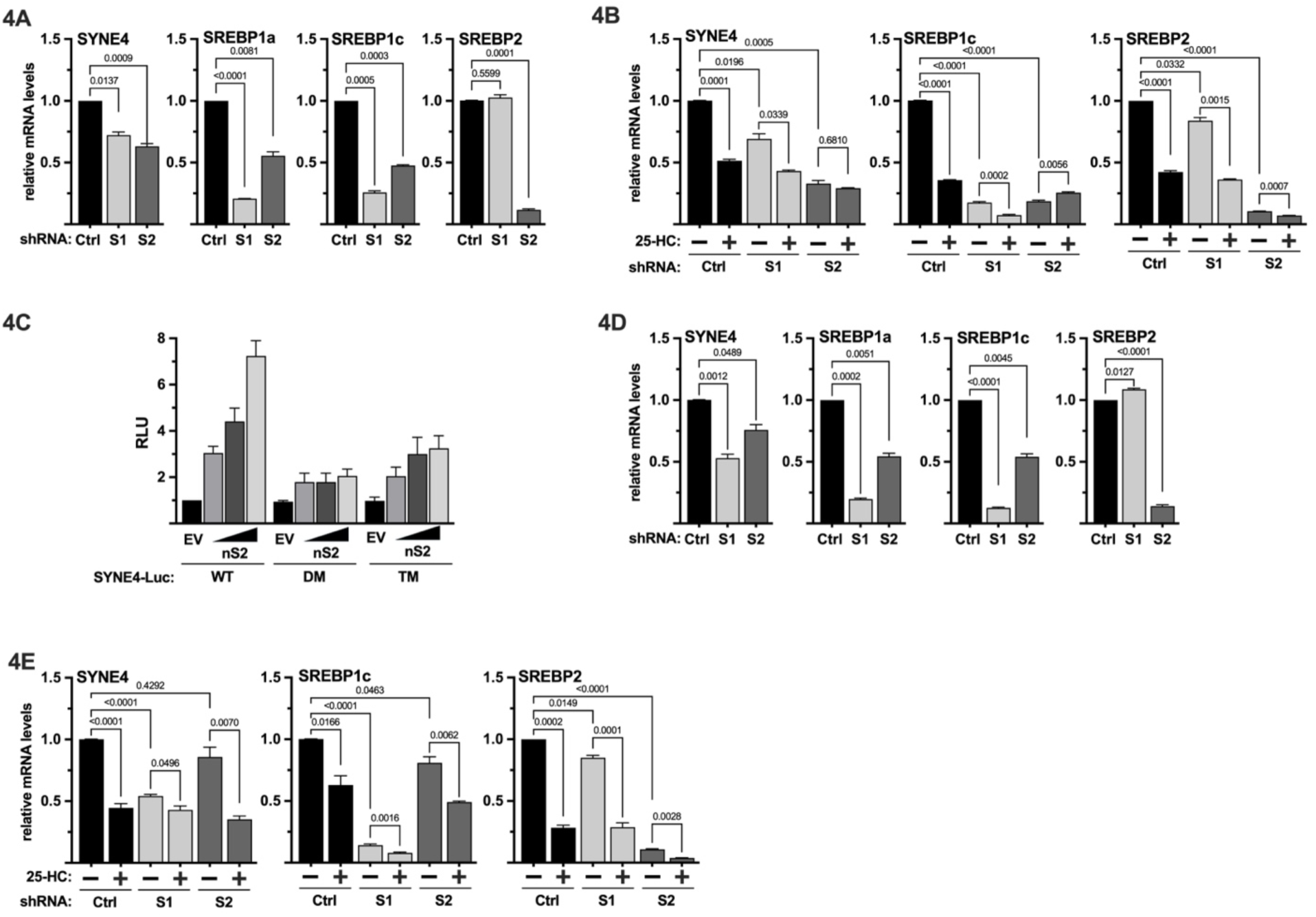
The expression of the SYNE4 is regulated by sterols and SREBP1/2. (**A**) HepG2 cells were transduced with lentiviruses expressing non-targeted (Ctrl), SREBP1 (S1) or SREBP2 (S2) shRNA and left in regular growth media. Ninety-six hours following transduction, mRNA was extracted and used for qPCR with primers specific for SYNE4, SREBP1a, SREBP1c, and SREBP2. (**B**) HepG2 were transduced as in A. Ninety-six hours following transduction, the media was changed to lipoprotein-deficient media to activate SREBP1/2. Where indicated, the media was supplemented with 25-hydroxycholesterol (25-HC) to block the activation of SREBP1/2. Twenty-four hours following the addition of 25-HC, mRNA was extracted and used for qPCR with primers specific for SYNE4, SREBP1c, and SREBP2. (**C**) HepG2 cells were transfected with SYNE4 promoter reporter constructs, either wild-type (WT), DM or TM, together with an empty expression vector (EV) or increasing amounts of nuclear SREBP2 (nS2). Forty-eight hours after transfection, the cells were lysed, and luciferase activity was measured. The cells were also transfected with the β-galactosidase gene as an internal control for transfection efficiency. Luciferase values (relative light units, RLU) were calculated by dividing the luciferase activity by the β-galactosidase activity. (**D**) MCF7 cells were transduced with lentiviruses expressing non-targeted, SREBP1 (S1) or SREBP2 (S2) shRNA and left in regular growth media. Ninety-six hours following transduction, mRNA was extracted and used for qPCR with primers specific for SYNE4, SREBP1a, SREBP1c, and SREBP2. (**E**) MCF7 were transduced as in D. Ninety-six hours following transduction, the media was changed to lipoprotein-deficient media to activate SREBP1/2. Where indicated, the media was supplemented with 25-hydroxycholesterol (25-HC) to block the activation of SREBP1/2. Twenty-four hours following the addition of 25-HC, mRNA was extracted and used for qPCR with primers specific for SYNE4, SREBP1c, and SREBP2.

We have previously demonstrated that nesprin-4 affects the migration and invasion of breast cancer cell lines, including MCF7 cells. To determine if the expression of SYNE4 was regulated by SREBP1/2 in these cells, the experiment illustrated in Figure 4B was repeated in MCF7 cells. Again, the expression of SYNE4 was reduced in response to inactivation of both SREBP1 and SREBP2 in cells grown in regular media (Figure 4D). In contrast to HepG2 cells, inactivation of SREBP1 attenuated the sterol-dependent regulation of SYNE4, while inactivation of SREBP2 failed to do so (Fig. 4E), suggesting that the sterol-dependent regulation of SYNE4 is controlled by SREBP1/2 in a cell type dependent manner.

### The SREBP-SYNE4 signaling axis is bidirectional and nesprin-4 controls the expression and function of SREBP1c

To further explore the functional link between nesprin-4 and the SREBP pathway, we used lentiviral shRNA to knock down SYNE4 in HepG2 cells and monitored the expression of SREBP1/2 and some of their target genes. Surprisingly, we found that the expression of SREBP1c was significantly reduced in nesprin-4-deficient cells (Fig. 5A). Although we also observed a decrease in the expression of SREBP1a and SREBP2 in the nesprin-4-deficient cells, these changes were not of the same magnitude as that of SREBP1c (Fig. 5A). Interestingly, the expression of the fatty acid desaturase SCD1 and fatty acid synthase (FASN), classical SREBP1c target genes, was also significantly reduced in SYNE4 knockdown cells. Both FASN and SCD1 play important roles in lipid metabolism, with SCD1 desaturating the fatty acids produced by fatty acid synthase. In agreement with the limited effect on the expression of SREBP2, the inactivation of SYNE4 in HepG2 cells had limited effects on the expression of typical SREBP2 target genes involved in cholesterol synthesis and metabolism. The reduced expression of SREBP1 and SCD1 was also apparent at the protein level (Fig. 5B). The insulin-dependent expression and posttranscriptional activation of SREBP1c are important for organismal lipid metabolism. Interestingly, inactivation of SYNE4 in HepG2 cells reduced the insulin-dependent increase in active SREBP1 and SCD1 expression (Fig. 5C).

**Figure 5.**
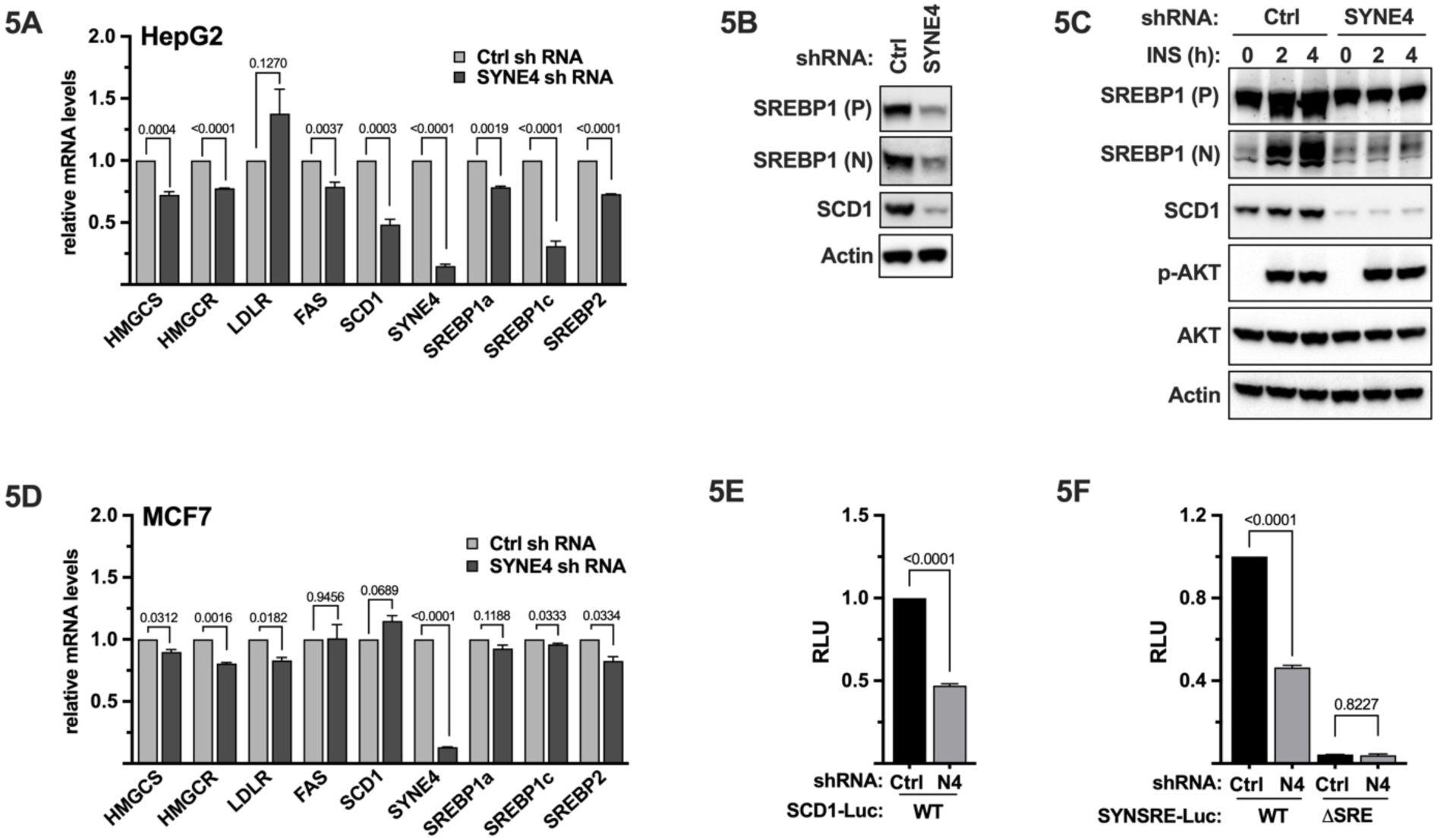
Nesprin-4 is a positive regulator of the SREBP pathway. (**A**) HepG2 cells were transduced with lentiviruses expressing non-targeted (Ctrl) or SYNE4 (N4) shRNA and left in regular growth media. Seventy-two hours following transduction, the media was changed to lipoprotein-deficient media to activate SREBP1/2. Twenty-four hours following the change of media, mRNA was extracted and used for qPCR with primers specific for SYNE4, SREBP1a, SREBP1c, SREBP1c, and the SREBP target genes HMG-CoA synthase (HMGCS), HMG-CoA reductase (HMGCR), fatty acid synthase (FAS), the LDL receptor (LDLr), and stearoyl-CoA desaturase (SCD1). (**B**) HepG2 cells were transduced as in A. Seventy-two hours following transduction, cells were placed in lipoprotein-deficient media overnight to activate SREBP1/2, followed by protein extraction. The extracted proteins were resolved on 4-12% SDS-PAGE gels, transferred to nitrocellulose membranes and probed with primary antibodies against SREBP1, SCD1, and β-actin. The SREBP1 antibody recognizes the two major forms of the protein, the precursor (P) and the transcriptionally active nuclear form (N). (**C**) HepG2 cells were transduced as in A. Seventy-two hours following transduction, cells were washed once in PBS and placed in serum-free media supplemented with 0.5% BSA overnight. The serum-starved cells were left untreated or treated with insulin for 2 or 4 hours, followed by protein extraction. The extracted proteins were resolved on 4-12% SDS-PAGE gels, transferred to nitrocellulose membranes and probed with primary antibodies against SREBP1, SCD1, AKT phosphorylated on serine 473 (P-AKT), total AKT, and β-actin. The SREBP1 antibody recognizes the two major forms of the protein, the precursor (P) and the transcriptionally active nuclear form (N). (**D**) MCF7 cells were transduced with lentiviruses expressing non-targeted (Ctrl) or SYNE4 (N4) shRNA and left in regular growth media. Seventy-two hours following transduction, the media was changed to lipoprotein-deficient media to activate SREBP1/2. Were indicated, the media was supplemented with 25-hydroxycholesterol (25-HC). Twenty-four hours following the change of media, mRNA was extracted and used for qPCR with primers specific for SYNE4, SREBP1a, SREBP1c, SREBP1c, and the SREBP target genes HMG-CoA synthase (HMGCS), HMG-CoA reductase (HMGCR), fatty acid synthase (FAS), the LDL receptor (LDLr), and stearoyl-CoA desaturase (SCD1). (**E**) HepG2 cells were transfected with a promoter reporter construct containing the sterol-responsive portion of the SCD1 promoter (SCD1-Luc) together non-targeted SYNE4 shRNA. Forty-eight hours following transfection, the media was replaced with lipoprotein-deficient media to activate SREBP1/2. Seventy-two hours following transfection, the cells were lysed, and luciferase activity was measured. (**F**). HepG2 cells were transfected with the HMG-CoA synthase promoter-reporter (SYNSRE-Luc), either wild-type (WT) or a version in which the SREBP binding site was deleted (ΔSRE). The cells in E and F were also transfected with the β-galactosidase gene as an internal control for transfection efficiency. Luciferase values (relative light units, RLU) were calculated by dividing the luciferase activity by the β-galactosidase activity.

The liver is the main site of de novo lipid synthesis and SREBP1/2 play important roles in these processes. To explore if the nesprin-4-dependent regulation of SREBP1c expression was conserved in MCF7 cells, the experiment illustrated in Fig. 5A was repeated in these cells. Although the expression of SREBP1c was slightly reduced in response to nesprin-4 inactivation, the effect was not as significant as that in HepG2 cells, and we did not observe any changes in the expression of FASN or SCD1 (Fig. 5D). However, the expression of SREBP2 and its target genes was slightly decreased in response to SYNE4 inactivation. Thus, the bidirectional SREBP-nesprin-4 signaling axis could play an especially important role in liver cells by regulating SREBP1c expression/function.

The expression of both SREBP1c and SCD1 is induced by SREBP1/2. However, these genes are also targets of other transcription factors. Therefore, we used promoter-reporter assays to explore if nesprin-4 affects the transcriptional activities of SREBP1/2. In these assays, we used reporter constructs in which the expression of the luciferase reporter gene was under the control the minimal sterol-responsive portions of the SCD1 and HMG-CoA synthase (HMGCS) promoters. These promoter-reporter constructs were transfected into HepG2 cells together with non-targeted or SYNE4 shRNA. As seen in Fig. 5E, inactivation of nesprin-4 reduced the activity of the SCD1 promoter, supporting our observations in nesprin-4 deficient cells. Inactivation of nesprin-4 also reduced the activity of the HMGCS promoter (Fig. 5F). Importantly, the nesprin-4 sensitivity of the HMGCS promoter was lost when the SREBP binding site in the promoter was deleted (Fig. 5F). These data suggest that nesprin-4 controls the activity of SREBP-dependent promoters through its effects on SREBP1/2 function, at least in HepG2 cells.

### Inactivation of nesprin-4 attenuates adipogenesis of adipose-derived stem cells

The expression of SREBP1c is highly induced during adipocyte differentiation [27–29], and has been shown to regulate both adipogenesis and lipid metabolism in mature adipocytes [27, 30, 31]. To determine if nesprin-4 affected the expression and function of SREBP1c during adipogenesis, human adipose-derived stem cells (ADSCs) were transduced with either non-targeted or SYNE4 shRNA. Ninety-six hours after transduction, the cells were either left untreated (undifferentiated) or differentiated towards mature adipocytes. The expression of SREBP1c, its target genes and established adipogenic markers were quantified by qPCR. As seen in Fig. 6A, the expression of SREBP1c and its target genes fatty acid synthase (FASN) and SCD1 were significantly reduced in the differentiated SYNE4-deficient cells. The same was true for all adipogenic markers analyzed (Fig. 6A). It is important to note that the inactivation of nesprin-4 failed to have any major effect on the expression of SREBP1a or SREBP2 (Fig. 6B), a result very similar to that observed in SYNE4-deficient HepG2 cells (Fig. 5A). The reduced adipogenic potential of SYNE4-knockdown ADSCs was confirmed when monitoring the accumulation of intracellular lipid droplets. Non-targeted ADSCs displayed an obvious increase in cells containing lipid droplets in response to adipogenic stimulation, while very few SYNE4-deficient ADSCs accumulated lipid droplets under the same conditions (Fig. 6C). Thus, the nesprin-4–SREBP1c axis is not specific for HepG2 cells or cancer cell lines but is also active in primary human cells. Importantly, nesprin-4 regulates adipogenesis, a biological process tightly associated with whole-body metabolism and SREBP1c activity.

**Figure 6.**
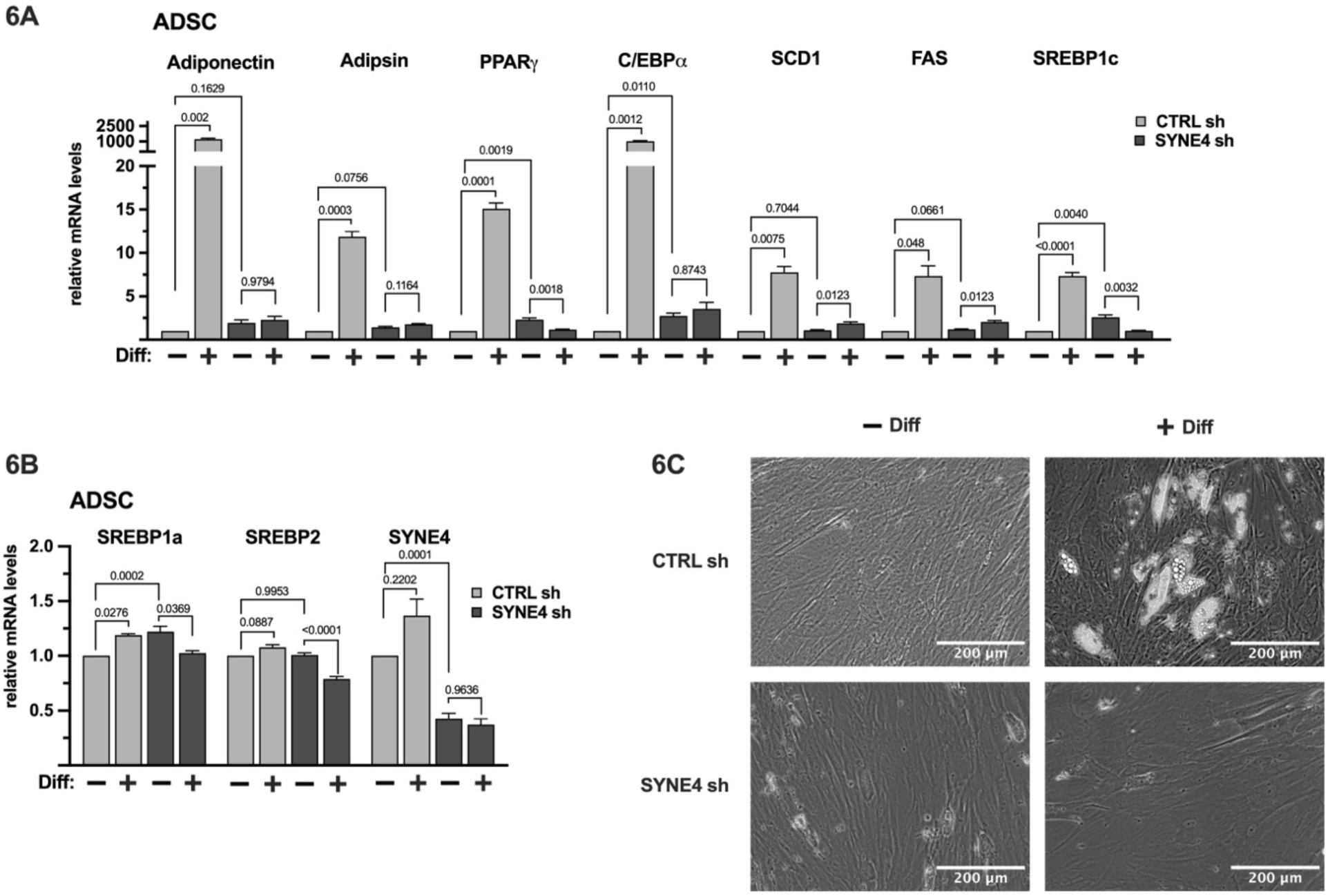
Nesprin-4 is required for adipogenesis in primary human ADSCs. (**A**) Human adipose-derived stem cells (ADSCs) were transduced with either non-targeted or SYNE4 shRNA. Ninety-six hours after transduction, the cells were either left untreated undifferentiated (–Dib) or differentiated towards mature adipocytes (+Diff). The expression of SREBP1c, its target genes and established adipogenic markers were quantified by qPCR. (B) ADSCs were transduced and treated as in (A). The expression of SREBP1a, SREBP2 and SYNE4 was quantified by qPCR. (**C**) ADSCs were transduced and treated as in (A). Photographs were captured on an EVOS M5000 Imaging System at 20x magnification.

Taken together, our data demonstrate that the expression of the LINC component nesprin-4 is controlled by members of the SREBP family of transcription factors, indicating a potential link between lipid metabolism and nuclear mechanotransduction. Importantly, we show that the functional interaction between SREBP1/2 and nesprin-4 is bidirectional in cell models characterized by high lipogenic potential, with nesprin-4 supporting SREBP1c expression and function. Thus, the interaction between lipid metabolism and the LINC complex, especially nesprin-4, could be seen as a therapeutic target in metabolic disease, which warrants further investigations of this novel functional interaction.

## Discussion

Here we establish a bidirectional link between the transcription factors SREBP1/2 and the LINC complex component nesprin-4. We demonstrate that the expression of nesprin-4, encoded by the SYNE4 gene, is regulated by intracellular sterol levels in HepG2 and MCF7 cells in a SREBP1/2-dependent manner. These observations are well aligned with the identification of potential SREBP binding sites in the SYNE4 promoter in genome-wide ChIP-sequencing studies performed as part of the ENCODE project, including in HepG2 and MCF7 cells (endcodeproject.org). Importantly, the SREBP binding sites identified in this report overlaps with those identified in these ChIP-sequencing studies. Using ATAC-sequencing, it has been suggested that the SYNE4 proximal promoter is located within an accessible/open region of chromosome 19 and that the promoter is associated with epigenetic marks typically found at active promoters, both in HepG2 and MCF7 cells (https://genome-euro.ucsc.edu). Notably, the accessible chromatin structure and the activating epigenetic marks found at the SYNE4 promoter in these two cell lines are retained in human primary liver and breast tissue (https://genome-euro.ucsc.edu). Taken together, these observations strongly support the notion that the SREBP-dependent SYNE4 promoter described in this report is functional in vivo, not only in cancer cell lines.

Although the basal expression of SYNE4 in MCF7 and HepG2 cells was dependent on both SREBP1 and SREBP2, the sterol-dependent regulation of the gene differed between the two cell lines. In HepG2 cell, SREBP2 was required for the sterol-dependent regulation of SYNE4, while SREBP1 was required for its sterol-dependent regulation in MCF7 cells. This observation demonstrates that different cell types can use different SREBP paralogs to regulate the expression of the same gene. This could be explained by differences in the expression of SREBP1 and SREBP2 in these two cell lines. The SREBP1a gene is derepressed in many tumors and cancer cell lines because of its ability to promote the expression of genes involved in both fatty acid/triglyceride and cholesterol biosynthesis, which would support the increased demand of membrane synthesis in cancer cells [32–34]. However, it is possible that the dominant role of SREBP2 in the control of sterol-dependent transcription has been retained in HepG2 cells because of the hepatic origin of this cell line. It will be interesting to determine if this type of paralog preference between cell and tissue types is also found in vivo.

The activation of SREBP1/2 is primarily controlled by the intracellular levels of cholesterol and in response to insulin signaling [35–38]. The precursor forms of SREBP1/2 reside in the ER membrane and are activated when the levels of cholesterol in the ER membrane fall below a certain threshold. The activation of SREBP1/2 requires the SCAP-dependent transport of the precursor molecules from the ER to the Golgi. Nesprin-4, like all members of the nesprin family, is a single-pass membrane protein that is associated with the outer nuclear membrane (ONM), which is continuous with the ER membrane. At the nuclear envelope, nesprin-4 forms part of the LINC complex that transmits mechanical cues from the cytoskeleton to the nucleus and chromatin by forming complexes with SUN1/2 embedded in the inner nuclear membrane (INM) [4, 6]. As far as we know, there have been no previous reports of a direct link between the SREBP1 pathway and the LINC complex. However, the nuclear form of SREBP1 has been found to interact with lamin A/C, an interaction that affects their transcriptional activity [39–41]. Interestingly, SUN1/2 physically interact with components of the nuclear lamina, including lamin A/C, as well as with chromatin [42–44]. Whether components of the LINC complex interact with components of the SREBP pathway is unknown, and this possibility should be explored in future work, especially since the SREBP precursor proteins have been reported to be associated with the nuclear membrane. It is also important to note that SREBP1 activation has been shown to respond to mechanical cues [45–47]. Notably, it has been demonstrated that SREBP1 is inactivated in cells grown on a stiff surface [45–47], a situation associated with increased mechanical strain on the nucleus. However, SREBP1 is activated in cells grown on a soft surface when the cell and nucleus experience less mechanical strain. Importantly, it has been demonstrated that the differentiation of mesenchymal stem cells to mature adipocytes, a process involving SREBP1c, is enhanced when cells are grown on a soft surface [46]. These observations point to a connection between the mechanotransduction machinery (potentially involving LINC complexes) and the SREBP mediated lipid synthesis.

Interestingly, several reports have linked the INM with lipid synthesis/metabolism [48–55]. The lipid profile of the INM is distinct from that of the ONM [48–55]. Notably, several enzymes involved in lipid synthesis have been found to associate with the INM. In addition, lipin-1, a lipid phosphatase involved in glycerolipid synthesis is known to shuttle between the cytoplasm and the INM, where it appears to control nuclear lipid droplet biogenesis [54, 56]. Of note, lipin-1 is both a target and regulator of SREBP1 [57, 58]. Nuclear lipid droplets have been identified and characterized in multiple cell types ranging from yeast to human neuronal cells [54, 56, 59, 60]. Nuclear lipid droplets are generated from the INM and may play an important role in controlling the lipid composition of the INM [54, 56, 59]. Indeed, an excess in saturated lipids in the INM is detrimental for nuclear integrity and function [61, 62]. Interestingly, SUN2 has the capacity to sense the lipid composition of the INM and detect lipid packing defects [63]. Lipid packing is controlled, at least in part, by the balance between saturated and unsaturated fatty acids within membrane glycerolipids. SUN2 has been shown to be degraded in response to changes in the ratio of saturated/unsaturated fatty acids in the INM [63]. Interestingly, we find that nesprin-4 supports the expression and function of SREBP1c, an important regulator of fatty acid synthesis and desaturation, especially in liver. Consequently, inactivation of nesprin-4 reduced the expression of SREBP1c target genes involved in fatty acid synthesis, including fatty acid synthase and SCD1. Thus, nesprin-4 could support SREBP1c function to maintain a critical balance between saturated and unsaturated fatty acids in the nuclear envelope, thereby preserving nuclear integrity. It is tempting to speculate that the degradation of SUN2 in response to changes in the saturated/unsaturated fatty acid ratio in the INM is sensed by nesprin-4, which in turn activates the SREBP1c-SCD1 axis. It is important to note that the inactivation of the SREBP target gene lipin-1, which also functions as a regulator of SREBP1, results in lipid packing defects and loss of nuclear integrity. Importantly, it has been demonstrated that lipin-1 loss affects the stability of SUN2. Thus, the SREBP pathway could potentially be an important regulator of nuclear functions/integrity by controlling the expression of SYNE4, SCD1 and lipin-1. This possibility will be an interesting focus of future investigation.

While we saw a significant relationship between SYNE4 knockdown and a decrease in SREBP1c and SCD1 expression in HepG2 cells, we did not observe the same in SYNE4 knockdown MCF7 cells. This could potentially be explained by a difference in the expression of SYNE4 and/or knockdown efficiency between these cells. If the basal expression of SYNE4 is higher in MCF7 cells compared to HepG2 cells, the remaining nesprin-4 protein may be sufficient to maintain SREBP1c expression/function. Alternatively, HepG2 cells are dependent on SCD1, while MCF7 rely on other fatty acid desaturases that are not under SREBP1c control. A third possibility is that the nesprin-4-mediated regulation of SREBP1c expression/function is specific for hepatocytes. However, this hypothesis does not align with our observation that the inactivation of nesprin-4 in adipose-derived stem cells (ADSCs) results in the loss of SREBP1c induction during adipogenesis without affecting the expression of SREBP1a or SREBP2. Importantly, inactivation of nesprin-4 in these primary progenitor cells significantly reduced their adipogenic potential.

Given the therapeutic potential for the nesprin-4-mediated control of SREBP1c, it will be important to determine the underlying mechanism(s) for how nesprin-4 promotes SREBP1c expression and activity. One possibility is that nesprin-4 physically interacts with the precursor form of SREBP1c, either in the ER or the ONM, and that this results in the proteolytic activation of SREBP1c. However, SREBP1/2 activation depends on the SCAP-mediated transfer of SREBP precursor molecules from the ER to the Golgi, where they are proteolytically activated [19, 20]. It seems unlikely that a component of the LINC complex is shuttled between these two organelles. Of note, a SCAP-independent mechanism to activate SREBP1c was recently described [64, 65]. This mechanism involves RHBDL4, an ER-resident protease that cleaves and activates SREBP1c. The authors demonstrate that the enzymatic activity of RHBDL4 is enhanced by saturated fatty acids and inhibited by polyunsaturated fatty acids [64], suggesting that the RHBDL4-dependent activation of SREBP1c provides cells with the possibility to sense and respond to changes in fatty acid saturation. Since nesprin-4 controls SCD1 expression, it will be interesting to determine if nesprin-4 plays a role in this fatty acid-controlled regulation of SREBP1c. It is interesting to note that the cytoplasmic domain of nesprin-4 contains a leucine-zipper domain [7]. The same domain is found in nuclear/active form of SREBP1/2 where it promotes transcription factor homo and heterodimerization. Thus, it could be that nesprin-4 interacts with active SREBP1c molecules and enhancing its nuclear import. Interestingly, the LINC complex has been found to promote the nuclear import of the transcription factor βcatenin [66, 67].

Another possibility is that nesprin-4 controls the expression of SREBP1c indirectly through the ability of LINC complexes to interact with chromatin. The nuclear lamina is mainly associated with heterochromatin, and although the expression of SREBP1c is regulated by histone K9 methylation (H3K9me) [68, 69], it has not been reported to be associated with heterochromatin. However, a recent study reported that the accessibility of the SREBP1 promoter region and the expression of SREBP1c was enhanced in Lamin A knockout oligodendrocytes [70]. Lamin A loss is associated with reduced levels of heterochromatic marks, including H3K9me3 and H3K27me3 [71, 72], suggesting that the SREBP1 gene could be associated with the nuclear lamina and/or heterochromatin. The genome-wide distribution of heterochromatin differs between cell types, which could explain the different responses to nesprin-4 loss between HepG2 and MCF7 cells. This possibility could be explored by mapping the epigenetic marks associated with the SREBP1 promoter in wild-type and nesprin-4-deficient HepG2 and MCF7 cells.

In liver cells, genes involved in fatty acid and triglyceride synthesis are controlled by SREBP1c, while genes involved in cholesterol synthesis and metabolism are controlled by SREBP2. The activation of SREBP2 is mainly controlled by intracellular cholesterol levels, while the activation of SREBP1c is insulin responsive [17, 19]. Paradoxically, insulin-dependent overactivation of SREBP1c in the liver is frequently seen in insulin resistant individuals, including T2D patients, which exacerbates the metabolic health of these individuals. The reason for this paradox is that these patients develop what is known as selective insulin resistance in the liver [73]. This means that insulin is unable to inhibit glucose output from the liver, while the insulin-dependent activation of SREBP1c is intact. As a result, lipid synthesis, especially the synthesis of triglycerides, increases. If untreated, the progressively increasing levels of circulating insulin seen in T2D patients result in an overproduction of triglycerides in the liver, resulting in hypertriglyceridemia and the development of non-alcoholic fatty liver disease. Hypertriglyceridemia can result in the deposition of fatty acids in peripheral tissues, including muscle and adipose tissue, thereby worsening the insulin resistance of these tissues. Thus, SREBP1c and the mechanisms controlling its activity are valid targets in metabolic disease, especially T2D. Interestingly, we found that the expression and insulin responsiveness of SREBP1 were attenuated in nesprin-4-deficient HepG2 cells, resulting in the reduced expression of SCD1. To the best of our knowledge, no SREBP1c-specific therapeutics have been developed/reported. Consequently, the nesprin-4-dependent regulation of SREBP1c expression/function could be considered a valid therapeutic target in metabolic disease. It will therefore be important to determine if nesprin-4 has an impact on the expression/function of SREBP1c in primary hepatocytes and/or in vivo and whether the inactivation of SYNE4/nesprin-4 influences SREBP1c activity in mouse models of insulin resistance.

In summary, this report identifies a bidirectional signaling axis that links nuclear mechanotransduction to the control of lipid metabolism. Initially, we identify the LINC component nesprin-4/SYNE4 as a novel SREBP target gene. Subsequently, we show that nesprin-4 promotes the expression and function of SREBP1c. SREBP1c is an important regulator of fatty acid and triglyceride synthesis, and we show that nesprin-4 loss reduces the expression of genes involved in fatty acid synthesis, especially the fatty acid desaturase SCD1. The balance between saturated and unsaturated fatty acids plays an important role in controlling membrane function and integrity, including in the inner and outer nuclear membrane. Importantly, SREBP1c and the mechanisms controlling its expression and/or function are valid targets in metabolic disease, including type 2 diabetes and obesity. Thus, the nesprin-4-dependent regulation of SREBP1c could become a therapeutic target for these diseases, warranting further work on this novel regulatory pathway.

## Material and Methods

### Cell Culture and Treatments

MCF7 (HTB-22), HepG2-C3A (CRL-10741), HeLa (CCL-2) and HEK293 (CRL-1573) cells were obtained from American Type Cell Culture Collection (ATCC). Cell culture media, supplements and reagents were from Gibco. HepG2-C3A cells were cultured in MEM media supplemented with 10% FBS, non-essential amino acids, sodium pyruvate, Glutamax and antibiotic-antimycotic, and MCF7, HeLa, and HEK293 cells were cultured in DMEM media supplemented with 10% FBS in addition to the supplements mentioned above. Where indicated, HepG2 and MCF7 cells were grown in media in which FBS was replaced by lipoprotein-deficient sera (Sigma-Merck) to promote the activation of SREBP1/2. Human adipose-derived stem cells (ADSCs) were obtained from ThermoFisher (R7788115) and cultivated in MesenPRO RS Medium (ThermoFisher, 12746012) and differentiated using the StemPro Adipogenesis Differentiation Kit (ThermoFisher, A1007001) as described previously [74].

### Plasmid DNA

The lentiviral shRNA constructs targeting human SREBP1 and SREBP2 were from Horizon Discovery and have been described previously [75–77]. The lentiviral shRNA constructs targeting human SYNE4 (VB900137-6002rva and VB900137-6003bzy) were purchased from VectorBuilder. The pGL2-SYNSRE-luciferase, pGL2-SYNSREι1SRE-luciferase and SCD1-Luc promoter-reporter constructs and the expression vectors for nuclear SREBP1a, SREBP1c and SREBP2 have been described previously [77–81].

### SYNE4 promoter cloning

Genomic DNA was extracted from MCF7 cells using the DNeasy Blood & Tissue Kit (QIAGEN) according to the manufacturer’s instructions. and the Nesprin-4 promoter was PCR amplified using the following primers incorporating restriction sites for KpnI and HindIII; KpnI-Nesp4-F CGGCCGGTACCCCGTGGGCAAAGGGGGCTCCCCTGG, HindIII-Nesp4-R GCCAAGCTTGGCTGGGGGCCTGGGGACACAAAGTCAGG. The PCR cycling conditions included an initial denaturation at 95°C for 2 minutes, followed by 35 cycles of 95°C for 30 seconds, 66°C for 30 seconds, and 72°C for 1 minute, with a final extension at 72°C for 7 minutes. After amplification, the PCR products were analyzed by running them on a 1% agarose gel to verify the size of the product. The PCR product was purified using the QIAquick PCR Purification Kit (QIAGEN #28104) according to the manufacturer’s protocol.

Subsequently, 1µg of the purified PCR product and 1µg of the vector were digested with high-fidelity KpnI and HindIII restriction enzymes for 2 hours at 37°C. The cleaved DNA was subsequently ligated into pGL4.12 (Promega).

### Q5-Site Directed Mutagenesis

Site-directed mutagenesis was performed via PCR using specific primers designed using the online NEB primer design software (https://nebasechanger.neb.com/). PCR amplification was carried out using Q5 Master Mix (NEB #E0554, BioLabs) in a 25 µL reaction mixture, which contained 12.5 µL of 2x Q5 Master Mix, 1.25 µL of 10 µM forward primer, 1.25 µL of 10 µM reverse primer, 9 µL of H2O, and 1 µL of 25 ng/µL template DNA. The PCR cycling conditions included an initial denaturation at 98°C for 30 seconds, followed by 25 cycles of 98°C for 10 seconds, 60°C for 30 seconds, and 72°C for 3 minutes, with a final extension at 72°C for 2 minutes. After PCR amplification, the product was mixed with 2x KLD (Kinase, Ligase, and DpnI) reaction buffer, KLD enzyme mix, and nuclease-free water in a total volume of 10 µL for one reaction. The reaction mixture was incubated at room temperature for 5 minutes to facilitate the ligation of the PCR product into the vector. For the transformation, the KLD reaction was added to 50 µL of competent E. coli cells. After incubating on ice for 30 minutes, the cells were subjected to heat shock at 42°C for 30 seconds. The transformed cells were plated onto LB agar plates containing ampicillin and incubated overnight at 37°C. Individual colonies were picked from the plates and transferred to 3 mL of LB broth supplemented with 100 µg/mL ampicillin. The cultures were incubated overnight at 37°C with shaking to allow for bacterial growth and plasmid amplification. Plasmid DNA was then extracted and purified using the QIAprep Spin Miniprep Kit (Qiagen) according to the manufacturer’s instructions. The purified plasmid DNA was subsequently used for further analysis and digested with restriction enzymes.

### Sanger Sequencing

The mutant SYNE4 promoter-reporter constructs described above were sequenced to confirm the individual deletions. The sequencing process began with PCR amplification using the following two primers: RV primer 3 (RV3)-F CTAGCAAAATAGGCTGTCCC, GL primer 2 (GL2)-R CTTTATGTTTTTGGCGTCTTCCA. The PCR master mix, with a total volume of 10 µl per reaction, consisted of 1.5 µl of BigDye Terminator v3.1, 2 µl of 5x Sequencing Buffer, 1.0 µl of 3 µM primer, 1.0 µl of DNA template (100-150 ng), and 4.5 µl of nuclease-free water. After amplification, the samples were purified using a master mix containing 45 µl of SAM TM Solution (ThermoFisher, MA01730) and 10 µl of X Terminator Solution Buffer (ThermoFisher, MA01730), yielding a total volume of 55 µl per reaction. Fifty-five µl of this master mix was added to each PCR tube, vortexed for 20 minutes at room temperature, and centrifuged at 1000xg for 2 minutes at 4°C. Finally, 15 µl of the supernatant was transferred into the sequencing plate for analysis. The resulting sequencing chromatograms were visualized and analyzed using SnapGene software to assess sequence quality and confirm base-calling accuracy.

### Lentivirus Production and Transduction

HEK293 cells grown in 10 cm dishes were used to produce all lentiviruses. Twelve μg of lentiviral DNA was co-transfected with 15 μl Trans-Lentiviral shRNA Packaging Kit (Horizon Discovery, TLP5912) by the calcium phosphate precipitation transfection method. Forty-eight hours after transfection, media was collected and filtered through 0.45 μm syringe filters and the viruses were stored in aliquots at –80°C. Target cells were transduced in regular media containing 8μg/ml polybrene for 16 hours, followed by 3 to 4 days of puromycin selection (2 μg/ml). The viruses expressing shRNA targeting human SREBP1, SREBP2, and nesprin-4 were produced and used as pools consisting of two to three shRNAs.

### Antibodies and Reagents

Antibodies against SCD1 (2438S), phosphorylated AKT (4058S), and AKT (4691S) were purchased from Cell Signaling Technology. Antibodies against SREBP1 (sc-8984 and sc-13551) were purchased from Santa Cruz Biotechnology, and the β-actin antibody (A5441) was purchased from Sigma-Merck. Horseradish peroxidase (HRP)-conjugated anti-rabbit IgG (G21234) and anti-mouse IgG (62-6520) antibodies were purchased from Invitrogen. Chemicals were obtained from Sigma-Merck, unless otherwise indicated.

### Cell lysis and immunoblotting

Cells were lysed in buffer A (50 mM HEPES (pH 7.2), 150 mM NaCl, 1 mM EDTA, 20 mM NaF, 2 mM sodium orthovanadate, 10 mM β-glycerophosphate, 1% (w/v) Triton X-100, 10% (w/v) glycerol, 1 mM phenylmethylsulfonyl fluoride (PMSF), 10 mM sodium butyrate, 1% aprotinin, 0.1% sodium dodecyl sulfate (SDS), and 0.5% sodium deoxy-cholate (DOC)) and cleared by centrifugation. Proteins were resolved by SDS–PAGE (4–12% Bis-Tris; Invitrogen) and transferred to nitrocellulose membranes (Cytiva). Membranes were blocked in 5% BSA in PBS containing 0.05% Triton X-100, probed with primary and HRP-conjugated secondary antibodies, and visualized by chemiluminescence on an iBright CL1500 (Invitrogen).

### Protein Purification

Cultures of E. coli (BL21) transformed with expression vectors for 6xHis nSREBP1a were induced with IPTG (0.75 mM) and incubated overnight at room temperature with shaking to allow protein expression. Cells were harvested by centrifugation, and the cell pellets resuspended in 20 ml PBS (ice-cold) containing protease inhibitors (PMSF and aprotinin) and sonicated on ice. Following sonication, Triton-X-100 was added to a final concentration of 1% and the suspension was kept in an end-over-end mixer for 30 minutes at 4°C. The solubilized material was centrifuged at 10,000 rpm for 20 minutes at 4°C, and the supernatant was collected in new tubes. Clarified lysates were used to purify the HisSREBP1a using Ni-NTA (Millipore, P661) employing standard protocols [79]. The purified 6xHis-SREBP1a protein was eluted from the Ni-NTA beads with imidazole. All eluted proteins were dialyzed against PBS containing 20% glycerol and protease inhibitors (PMSF and aprotinin), aliquoted and stored at –80°C.

### Electromobility shift assays

Electromobility shift assays were performed as described earlier [79, 82], with the exception that unlabeled DNA probes were used. In brief, unlabeled DNA probes corresponding to the wild-type was generated by PCR using the pGL4-SYNE4 template. The primer sequences were as follows, CTA GCA AAA TAG GCT GTC CC and CTT TAT GTT TTT GGC GTC TTC CA. The 10x binding buffer contains 200 mM Tris-HCl pH 7.5 and 500 mM NaCl. The final reaction mixture contained 2 μl 10x binding buffer, 20% glycerol, 1 μg sheared salmon sperm DNA (non-competitive DNA), 1 mM MgCl2, 1 mM dithiothreitol, and 0.5 μg bovine serum albumin (BSA). Five hundred ng of DNA probe were incubated in the absence or presence of 6-His-SREBP1a. In the experiment illustrated in Figure 1C, recombinant SREBP1a were replaced with nuclear extracts from HeLa cells grown in lipoprotein-deficient media to induce SREBP1/2 activation, in the absence or presence of an SREBP1 antibody (0.5 μg). The reactions were separated on 4% polyacrylamide gels with 0.5x TBE buffer, stained with SYBR Safe, and visualized on an iBright CL1500 Imaging System (Invitrogen).

### Luciferase and β-galactosidase assays

Promoter-reporter assays were performed as described previously [75, 76, 78, 83]. In brief, HepG2 cells were transiently transfected with the indicated promoter-reporter genes in the absence or presence of the indicated expression and/or shRNA vectors. Luciferase activities were determined in duplicate samples as described by the manufacturer (Promega). Cells were also transfected with the β-galactosidase gene as an internal control for transfection efficiency. Luciferase values (relative light units, RLUs) were calculated by dividing the luciferase activity by the β-galactosidase activity. The data represent the average −/+ SEM of at least three independent experiments performed in duplicates.

### RNA extraction and qPCR

RNA was extracted using GeneJet RNA Purification Kit (ThermoFisher). cDNA was generated using Applied Biosystems High-Capacity cDNA Reverse Transcription Kit. For qPCR, PowerUp SYBR Green Master Mix was used (Applied Biosystems), using hypoxanthine phosphoribosyltransferase (HPRT) as reference/housekeeping gene. HPRT1 The human primer sequences used to amplify target genes are provided in Table 1.

**Table 1.**
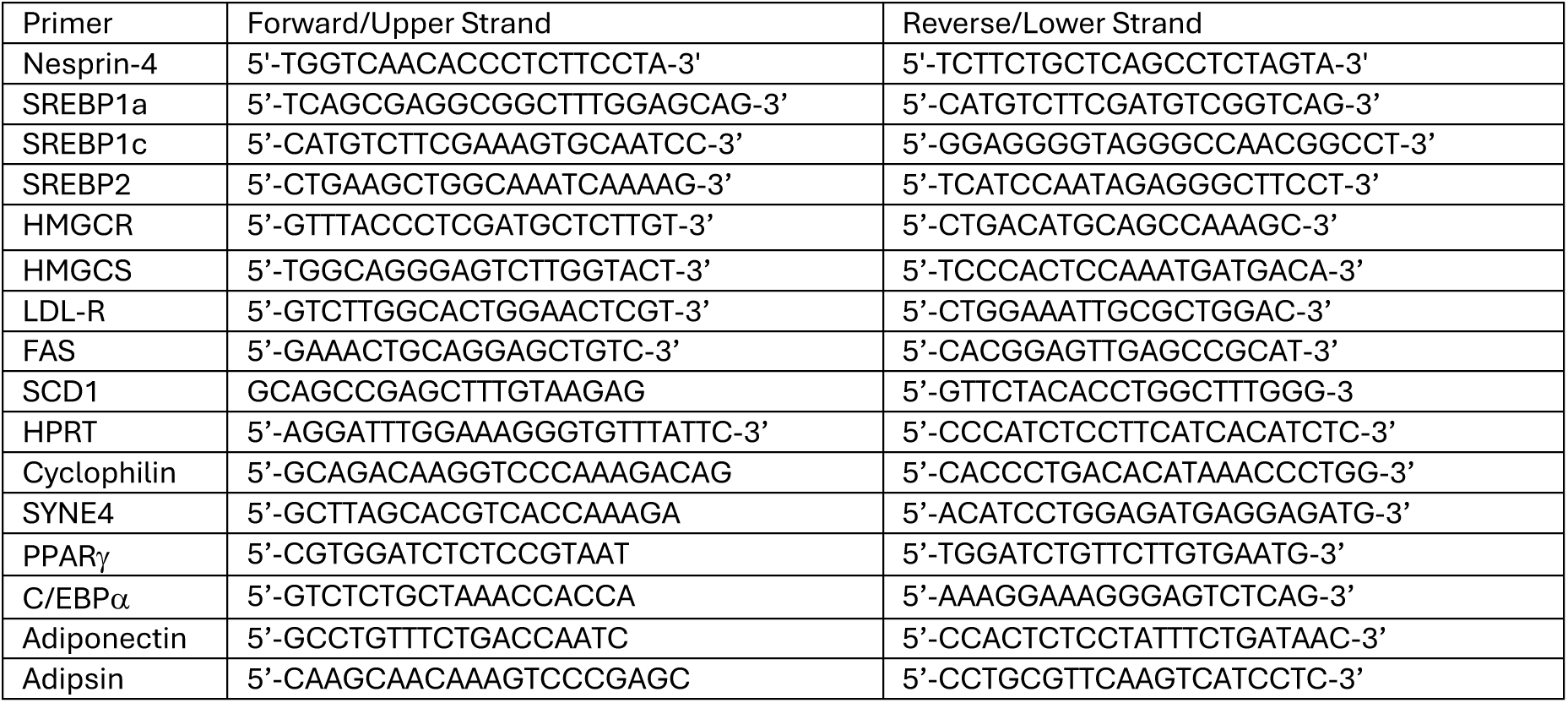
Human primer sequences used in the RT-qPCR assays.

### Statistical analysis

Statistical analyses were performed using GraphPad Prism version 10 (GraphPad Software). For comparisons involving more than two groups of a single experimental factor, Welch’s one-way ANOVA (Brown–Forsythe and Welch ANOVA test) was applied, as this approach does not assume equal variances. When the ANOVA indicated significance, Dunnett’s T3 multiple comparisons test was used for post-hoc pairwise group comparisons. For direct comparisons between two groups, Welch’s t-test was employed. Data are presented as mean ± SEM, unless otherwise indicated. A p-value < 0.05 was considered statistically significant.

