## Supplemental Figure 1 for "A bidirectional interaction between the SREBP pathway and the LINC complex component nesprin-4 controls lipid metabolism"

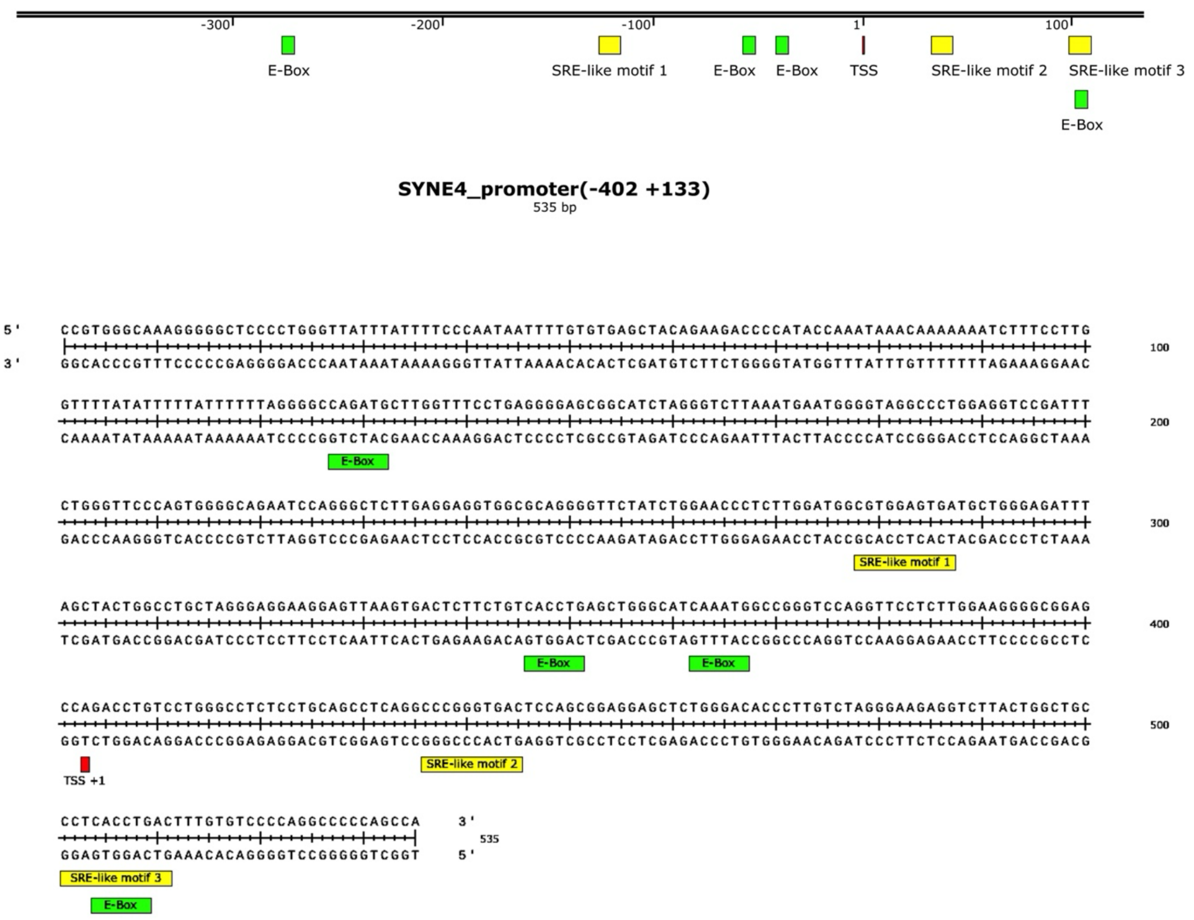


**Figure S1. The human SYNE4 promoter contains potential SREBP binding sites**. Illustration and sequence of the human SYNE4 promoter, including the location of E-box and potential SREBP binding (SRE-like) motifs and transcriptional start site (TSS).
